# Proteolytic dissection of eIF4G reveals the closed-loop mRNP as an architecture for translation repression

**DOI:** 10.64898/2026.04.06.716749

**Authors:** Ryan Johnston, Mollie A. Brekker, Noelle Khalil, Monty E. Goldstein, Anne Aldrich, Autumn O. Grimins, Sami Gritli, Assen Marintchev, Michael D. Blower, Mohsan Saeed, Shawn M. Lyons

## Abstract

Formation of a “closed-loop” mRNP, in which the 5′ cap and 3′ poly(A) tail are bridged by eIF4E-eIF4G-PABP interactions, has long been proposed to drive efficient translation initiation. Direct tests of this model in mammalian cells have remained elusive. Using auxin-inducible degron (AID) technology to acutely deplete eIF4G1, we find that global translation is only partially reduced and recovers without restoration of eIF4G1 levels. We identify eIF4G3 as an underappreciated contributor to basal translation that buffers translational output upon eIF4G1 loss without increased protein expression, explaining the modest defects observed in prior RNAi-based studies. Systematic replacement of eIF4G1 with defined cleavage products and interaction mutants reveals that PABP binding by eIF4G1 is dispensable for bulk translation initiation: the central caspase-3 cleavage fragment of eIF4G1 (casp3-cp^M^), which lacks the PABP-interaction domain, fully rescues global protein synthesis, and acute depletion of both major cytoplasmic PABP paralogs primarily destabilizes mRNAs rather than impairing initiation. In contrast, the N-terminal enteroviral 2A cleavage product (2A-cp^N^) is a potent, dominant translational repressor that requires simultaneous eIF4E and PABP engagement to form a dead-end closed-loop mRNP that sequesters initiation factors without enabling 43S recruitment. These findings reveal that the eIF4G-PABP closed-loop architecture is not required for productive initiation but can be actively co-opted for translational silencing. This explains why viral eIF4G cleavage, but not factor depletion, produces near-complete translational shutoff. The modular architecture of eIF4G enables diametrically opposing translational outcomes through selective proteolytic processing.

## Introduction

Cap-dependent translation initiation relies on the eukaryotic initiation factor 4F (eIF4F) complex, composed of the cap-binding protein eIF4E, the ATP-dependent RNA helicase eIF4A, and the large scaffolding protein eIF4G^1^. eIF4F mediates recruitment of the 43S pre-initiation complex (PIC) to the 5′ end of an mRNA, forming the 48S PIC, which includes the 40S ribosomal subunit, eIF3, and additional initiation factors such as eIF1, eIF2, and eIF5. eIF4G plays a central role in the assembly of this complex, coordinating interactions with eIF4E and PABP through its N-terminal domain, and recruiting eIF4A and eIF3 through its central MIF4G domain (**Figure 1A**).

**Figure 1.**
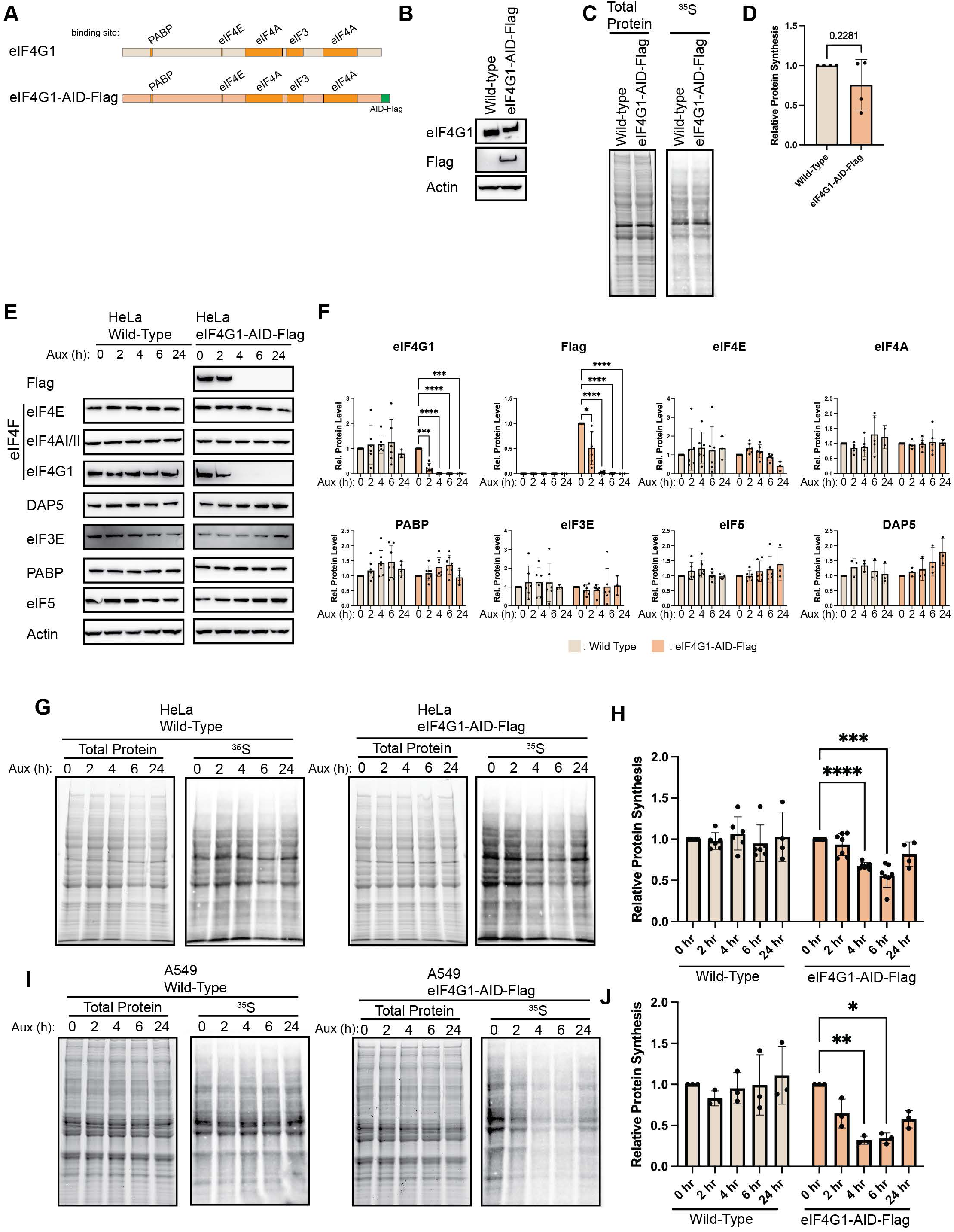
Basal translation is buffered upon acute loss of eIF4G1. (A) Schematic of eIF4G1 with relevant protein interaction domains indicated and indicating addition of AID-Flag tag to scale.(B) Western blotting of wild type or eIF4G1-AID-Flag cells demonstrates slight mobility shift in eIF4G1 and reactivity with Flag antibodies. (C – D) Basal translation is unaffected by tagging of eIF4G. (E & F) Addition of auxin results in rapid depletion of eIF4G1 as determined by eIF4G and Flag antibodies. Other interactors of eIF4G1 or associated eukaryotic initiation factors are unaltered upon eIF4G1 loss. Statistical significance determined by two-way ANOVA. (G & H) Autoradiography of HeLa eIF4G1-AID-Flag and wild type cells following auxin treatment at indicated times. Auxin has no effect on protein synthesis in wild type cells. Concurrent with depletion of eIF4G, protein synthesis rates decrease at 4 and 6 hours post-auxin treatment; however, these recover by 24 hours. (I & J) Autoradiography in A549 eIF4G1-AID-Flag cells confirms buffering capability upon chronic loss of eIF4G. Statistical significance determined by paired student’s t-test (D) or two-way ANOVA with Geisser-Greenhouse correction and Dunnett’s Multiple comparisons test with individual variances (F, H, J). Mean ± SD, n≥3. *p<0.0332, **p<0.0021, ***<0.0002, ****p<0.0001.

The interaction between eIF4G and PABP has long been proposed to promote the formation of a “closed-loop” mRNP, in which the 5’ cap and 3’ poly(A) tail are bridged through an eIF4E-eIF4G-PABP interaction^2^. The architecture has been proposed to enhance translational efficiency through promoting ribosome recycling and stabilizing initiation complexes at the 5′ end. This model was initially supported by genetic and biochemical studies in yeast showing that disruption of the eIF4G-PABP interaction impairs translation output^3,4^, and by *in vitro* reconstitution experiments demonstrating that PABP stimulates cap-dependent translation. However, whether these effects reflect direct kinetic enhancement of initiation or indirect stabilization of mRNA has been difficult to resolve, particularly *in vivo*. A key challenge has been that depletion of PABP produces confounding mRNA stability defects that cannot easily be separated from direct effects on initiation, and the extended timescales required for siRNA-based knockdown of eIF4G permit compensatory responses that obscure the acute translational requirement.

Because of its essential role in 48S PIC formation, eIF4G is a frequent target of translational control. Under growth-limiting conditions or cellular stress, hypophosphorylation of eIF4E-binding proteins (4EBPs) promotes high-affinity binding of 4EBPs to eIF4E, thereby preventing eIF4G association with the cap and inhibiting eIF4F-dependent initiation^5,6^. In response to various stresses ^7,8^ and during initiation of apoptosis^9^, caspase-3 (casp3) is activated and targets eIF4G for cleavage at two separate sites^10,11^. This distributes interaction domains among the resulting cleavage products. However, whether casp3 proteolytic cleavage of eIF4G simply disables translation initiation or generates functionally distinct regulatory fragments remains unclear; although, it has been shown that the middle fragment of eIF4G, which contains the MIF4G domain, associates with ribosomes^12^. This question is particularly relevant given evidence that sublethal casp3 activation can promote cell survival rather than apoptosis^7,8,13^, raising the possibility that eIF4G cleavage contributes to a protective translational response rather than simply silencing protein synthesis.

Similarly, enteroviruses produce a protease, known as 2A protease (2A^Pro^), that targets eIF4G at a single site that is distinct from casp3^14^. 2A^Pro^-mediated eIF4G cleavage separates the PABP and eIF4E binding domains from the eIF3 and eIF4A binding domains. Most studies evaluating enterovirus-mediated eIF4G cleavage have used poliovirus as model, but this feature is common among enteroviruses, including rhinoviruses and coxsackieviruses^15,16^. 2A^Pro^-mediated eIF4G cleavage is thought to promote viral mRNA translation by suppressing cap-dependent translation and biasing translation towards cap-independent modes of initiation. Enteroviral mRNAs contain internal ribosome entry sites (IRESes) that dispense for the need for eIF4E for initiation^17,18^. Instead, the C-terminal cleavage product of eIF4G (2A-cp^C^) is thought to bind to the IRES and recruit the 43S PIC^19–21^. However, the functional significance of the N-terminal cleavage product (2A-cp^N^), which retains the eIF4E- and PABP-binding domains of eIF4G but lacks the eIF3 and eIF4A interactions, has not been investigated.

eIF4G, like other components of the eIF4F complex, is encoded by multiple paralogs. While some paralogs appear to perform specialized functions, others may act redundantly with the canonical subunits; however, this possibility has not been directly tested. Consistent with this idea, several eIF4F components possess paralogs with divergent or context-specific roles. For example, eIF4E exists as 3 distinct paralogs: eIF4E (or eIF4E1), eIF4E2 (or 4EHP) and eIF4E3 ^22^. eIF4E2 primarily functions as a translational repressor^23,24^, although it can promote translation in specific contexts ^25,26^, whereas considerably less is known about eIF4E3, which has been implicated in the translation of specific mRNAs^27,28^. Similarly, eIF4A2 is highly homologous to eIF4A1^29^, but its cellular role has been controversial: while several studies suggest it can promote translation^30,31^, others indicate it can function as a translational repressor in specific contexts^32–34^. In contrast, eIF4A3 does not function in translation and instead acts as a core component of the exon junction complex involved in splicing ^35^.

Notably, eIF4G exists as two cap-dependent paralogs in mammals, eIF4G1 and eIF4G3 (eIF4GII) and a 3^rd^ paralog that does not interact with eIF4E known as eIF4G2 (DAP5). Despite its inability to bind eIF4E, DAP5 has been implicated in non-canonical modes of translation initiation ^36–40^. In contrast, physiological roles for eIF4G3 have been poorly defined ^41^. Evidence from poliovirus infection suggests that cleavage of eIF4G3 follows that of eIF4G1 and correlates with more complete inhibition of host protein synthesis ^42^, raising the possibility that eIF4G3 sustains translation when eIF4G1 activity is impaired. However, other studies have proposed that eIF4G3 instead performs specialized functions, such as regulating meiotic translation^43^ or promoting translation of hypoxia-responsive mRNAs ^26^. Whether eIF4G3 primarily functions as a backup for basal translation or supports selective translational programs therefore remains unresolved. This ambiguity is further compounded by the fact that eIF4G paralogs are differentially targeted during stress and viral infection^42,44–46^, suggesting that their functional relationships may only become apparent under conditions that perturb canonical eIF4F activity.

In this study, we engineered cells to leverage auxin-inducible degron (AID) technology to acutely deplete eIF4G1 and examine how translation initiation adapts to its loss. Using this approach, we identify eIF4G3 as a previously underappreciated contributor to basal protein synthesis that buffers translational output when eIF4G1 is compromised. We further show that depletion of eIF4G is mechanistically distinct from its proteolytic cleavage. Whereas enteroviral 2A protease-generated fragments act as dominant inhibitors of translation, caspase-3 cleavage products retain the ability to support protein synthesis. Together, these findings demonstrate that bulk translation initiation proceeds on linear mRNPs, independently of PABP-eIF4G bridging, and that the closed-loop architecture is instead co-opted as a mechanism of translational silencing during enterovirus infection. They further reveal that eIF4G proteolysis does not simply mimic factor loss but actively rewires translational control, with enteroviral 2A-gerated fragments acting as dominant repressors while caspase-3 cleavage products retain the capacity to support protein synthesis.

## RESULTS

### Generation of an auxin inducible degrons for eIF4G

Protein synthesis is an essential cellular process, which makes CRISPR-mediated knockout of many key initiation factors unfeasible. As a result, in mammalian systems, siRNA-mediated knockdown of individual eIFs has often been used to study their function. However, achieving robust depletion typically requires multi-day siRNA treatments, and these extended timescales permit secondary and compensatory cellular responses to the loss of essential initiation factors. To overcome this complication, we took advantage of the auxin-inducible degron (AID) system to study eIF4G function^47^. The *Arabidopsis thaliana* Auxin Signaling F-box 2 (AFB2) was inserted into AAVS1 locus using CRISPR-mediated homology directed repair. Into these cells, an AID and Flag tag was inserted at the 3’ end of the *eIF4G1* gene, which results in a C-terminally AID-Flag tagged protein upon expression (**Figure 1A - B**). Given eIF4G1’s role as a scaffolding protein, we worried that addition of tags to the endogenous eIF4G may have perturbed its activity. To rule out this possibility, we conducted [^35^S] metabolic labeling of wild-type or AID edited cells and found no difference between the two cells, establishing that editing does not alter global protein synthesis (**Figure 1C & D**). Thus, incorporating degron and epitope tags into the endogenous *eIF4G1* locus does not alter the functions of the eIF4G protein.

We next sought to determine the decay kinetics and the effect of basal translation on loss of eIF4G1. Wild-type or eIF4G1-AID-Flag cells were treated with 100 μM Auxin for 0, 2, 4, 6 and 24 hours. eIF4G was completely lost in the AID-tagged cell lines as monitored by western blotting to either eIF4G or to incorporated Flag epitopes within 4 hours (**Figure 1E & F**). Again, given eIF4G’s unique role as a scaffolding protein, we sought to assess whether destabilization of eIF4G1 co-depleted any of its binding partners. AID-mediated depletion has no statistically significant effect on the steady state levels of its direct interactors: eIF4E, eIF4AI/II, PABP or eIF3E. We also determined that eIF5, a component of the 43S PIC, were unchanged as was the eIF4G paralog, DAP5. These results demonstrate the efficacy of rapid targeted protein decay possible using AID-tagging. To determine the global effect of eIF4G depletion in HeLa cells, we monitored protein synthesis using [^35^S]-metabolic labeling. Cells were depleted of eIF4G1 as before but labeled with [^35^S]-Met/Cys for 15 minutes before harvesting, capturing nascent protein synthesis at that point (**Figure 1G & H**). Surprisingly, we found a decrease in protein synthesis upon depletion of eIF4G, but this reduction only reached ∼50% of basal protein synthesis upon 6 hours of depletion. These results are surprising in that eIF4G is known to be an essential factor in cap-dependent protein synthesis. Further, at 24 hours post-depletion, we observed a recovery in protein synthesis. This is despite no recovery of eIF4G1 as monitored by antibodies against eIF4G1 or incorporated Flag tag. To confirm these results and to establish the generality of this phenomenon, we generated a second eIF4G1-AID-Flag cell line in A549 cells (**Supplementary Figure 1**). As in HeLa cells, we found an initial reduction in translation, reaching its nadir by 6 hours, but partially recovering by 24 hours post-depletion (**Figure 1I & J**). While these results are contrary to what would be predicted based on the established role of eIF4G1, these results are largely in line with previous reports using siRNA-mediated depletion of eIF4G1 which demonstrated a 20% reduction in protein synthesis after 48 h of RNAi which recovered to baseline levels by 96 hours despite no increase in eIF4G1 levels^48^. These results suggest that basal translation does not require eIF4G or that it is buffered by additional cellular factors.

eIF4G1 is one of three eIF4G paralogs, the other two paralogous genes being eIF4G2 (DAP5) and eIF4G3 (eIF4GII). Historically, eIF4G2/DAP5 was thought to play a role in cap-independent translation initiation, as it lacks the eIF4E binding domain. However, recent work has demonstrated that eIF4G2/DAP5 is a key regulator in eIF3D-dependent translation initiation^49^ and is particularly relevant in metastatic potential of cells^37^. On the other hand, little is known about the biological role of eIF4G3. It shares approximately 48% similarity with eIF4G1 but retains known protein interaction domains with higher degrees of similarity (**Figure 2A**). eIF4G3 is incorporates into a non-canonical hypoxic eIF4F complex that specifically drives the translation of hypoxia-response genes, but little is known about its role in canonical translation initiation^50^. Previous work has shown that robust shutoff of host protein synthesis during poliovirus and rhinovirus infection correlates with eIF4G3 cleavage ^42,45^, suggesting that it does play a role in non-stressed protein synthesis.

**Figure 2.**
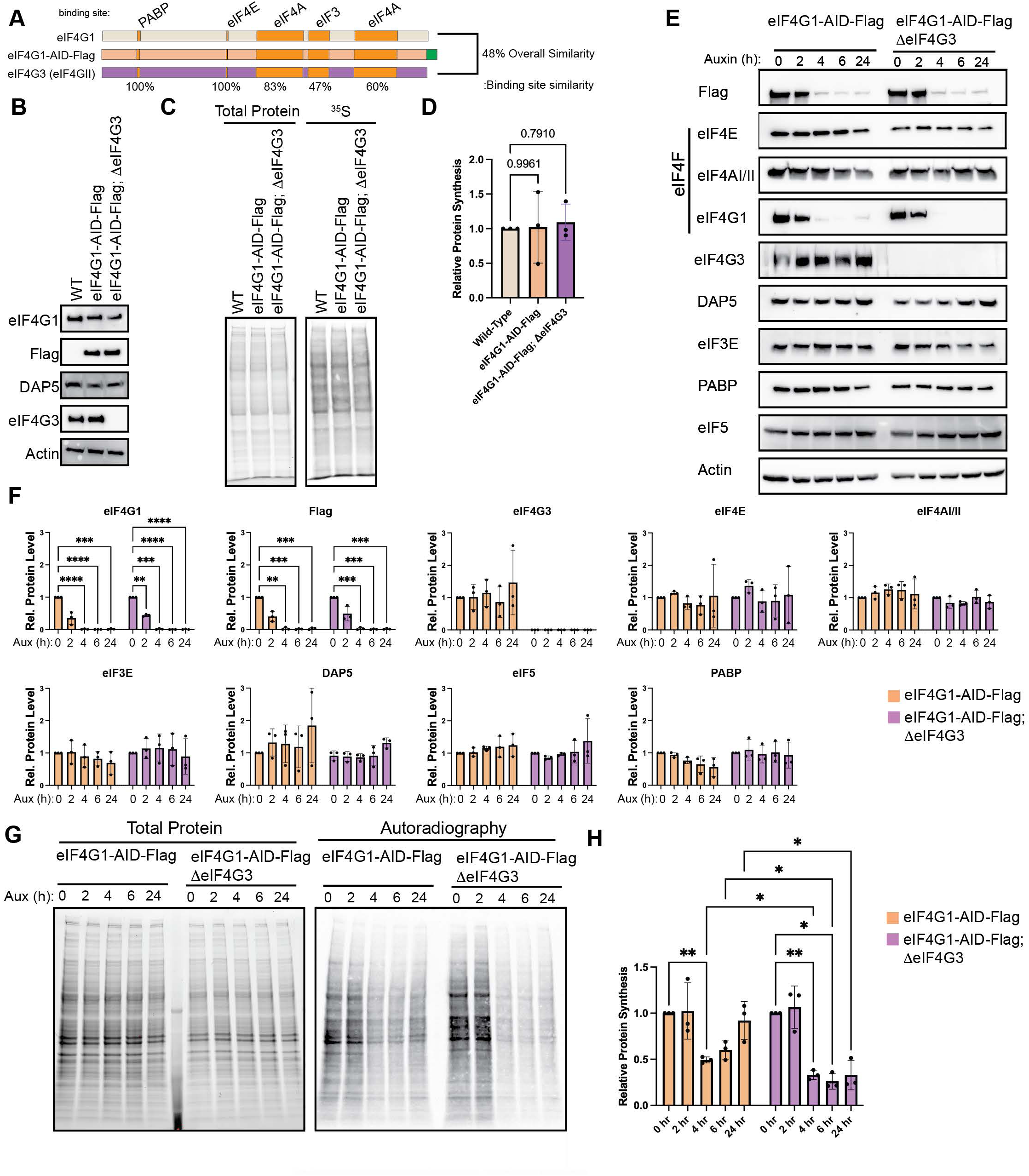
Loss of eIF4G3 ablates translational buffering upon chronic loss of eIF4G1. (A) eIF4G3 is a paralog of eIF4G1 which conserves all major protein:protein interaction domains. (B) CRISPR-mediated knockout of eIF4G3 does not alter the expression of other eIF4G paralogs. (C & D) Basal protein synthesis is unaltered in eIF4G3 knockout as determined by ^35^S incorporation. (E & F) Addition of auxin rapidly depletes eIF4G1 in ∆eIF4G3 background without co-depletion of other initiation factors. (G & H) Ablation of eIF4G3 abrogates translational buffering in response to eIF4G1 loss. Statistical significance determined by one-way ANOVA (D) or two-way ANOVA (F & H) with Geisser-Greenhouse correction and Dunnett’s Multiple comparisons test with individual variances. Mean ± SD, n≥3. *p<0.0332, **p<0.0021, ***<0.0002, ****p<0.0001.

Thus, we sought to test the role of eIF4G3 upon acute loss of eIF4G1 (4 - 6 hours post auxin) and during chronic loss when translational buffering has occurred (24 hours post auxin). To this end, we knocked out eIF4G3 using CRISPR/Cas9 (**Figure 2B**). Thus, upon treatment with auxin, these cells will express neither eIF4G nor eIF4G3. Critically constitutive loss of eIF4G3 has no effect on basal protein synthesis rates, indicating that eIF4G is sufficient to support global protein synthesis upon loss of eIF4G3 (**Figure 2C & D**). As before, auxin treatment resulted in a rapid depletion of eIF4G as monitored by flag and eIF4G antibodies and did not result in co-depletion of associated factors in eIF4G1-AID-Flag cells or eIF4G1-AID-Flag;∆eIF4G3 cells (**Figure 2E & F**). Further, we did not find an increase in eIF4G3 levels upon depletion of eIF4G1. This indicates that eIF4G3 was not responsible for stabilization of other eIFs upon loss of eIF4G1 and that their stability is independent of eIF4G1 or eIF4G3. We monitored global protein synthesis rates using [^35^S] metabolic labeling over the same time course (**Figure 2G & H**). This resulted in two important findings. First, knockout of eIF4G3 ablated translational buffering after auxin treatment. We found no recovery in protein synthesis when eIF4G1 was depleted. Second, the reduction in global translation was further enhanced by loss of eIF4G1 in a ∆eIF4G3 background. These results demonstrate that eIF4G3 is responsible for a portion of basal protein synthesis under homeostatic conditions. However, it suggests that a portion of eIF4G3 may be in a translational inactive state under normal conditions, and it can be “activated” upon loss or impairment of eIF4G1. This supposition is strengthened by the fact that we do not observe an increase in eIF4G3 protein levels during depletion.

We also monitored how knockout of eIF4G3 with and without eIF4G1 depletion affected cell growth (**Supplemental Figure 2**). In isolation, eIF4G3 knockout does not alter the rate of cell growth compared to either wild-type or eIF4G1-AID-Flag cells. Depletion of eIF4G1 with auxin abolishes net cell accumulation in both genetic backgrounds, with total cell numbers remaining stable over the course of the experiment. These data suggest that while eIF4G3 provides translational buffering in the absence of eIF4G1, the resulting translational program is insufficient to support robust cell expansion Despite these results, we still find observe 30 – 40% of global synthesis even upon loss of eIF4G1 and eIF4G3. Although this level of inhibition is significantly higher than found upon traditional siRNA-mediated depletion of eIF4G1, it is not the near complete inhibition of cellular protein synthesis seen during enterovirus infection upon eIF4G1 and eIF4G3 cleavage. However, we reasoned that depletion of eIF4G is not equivalent to protease mediated cleavage of eIF4G. Our system is uniquely positioned to ask how protease-generated cleavage products regulate cellular protein synthesis. To this end, we generated doxycycline (Dox)-inducible YFP-tagged constructs of eIF4G1 in eIF4G1-AID-Flag and eIF4G1-AID-Flag;∆eIF4G3 cells (**Figure 3A**). We generated dox-inducible YFP, full-length YFP-eIF4G1 or N-terminal (2A-cp^N^) and C-terminal (2A-cp^C^) cleavage products generated by enterovirus 2A proteases. Endogenous AID-tagged eIF4G1 was monitored using flag antibodies, while exogenous YFP tagged eIF4G1 fragments were monitored using anti-GFP antibodies, which also recognize YFP. Upon addition of auxin, endogenous eIF4G1 was completely ablated; but, upon addition of Aux and Dox, we rapidly replaced endogenous eIF4G1 with cleavage products in eIF4G1-AID-Flag (**Figure 3B**) and eIF4G1-AID-Flag;∆eIF4G3 (**Figure 3C**) cells. We tested the ability of eIF4G1 cleavage products to inhibit translation in eIF4G1-AID-Flag cells, in which eIF4G3 can buffer translation (**Figure 3D & E**), or in the absence of eIF4G3 (**Figure 3F & G**). As before, with eIF4G3 present (**Figure 3D & E**), basal protein synthesis levels are slightly reduced when eIF4G1 is lost (lanes 2, 5, 8, 11). YFP expression has no further effect and full length eIF4G1 slightly increases basal translation back to baseline levels. However, 2A-cp^N^, significantly inhibited translation, while 2A-cp^C^. Surprisingly, this effect was maintained even in the absence of eIF4G3 (**Figure 3F & G**). In these conditions, the rescue by full length eIF4G1 was more apparent due to the lack of eIF4G3 buffering. However, 2A-cp^N^ was still able to reduce the remaining protein synthesis by ∼50%. It is possible, although unlikely, that eIF4G1 that is undetectable by our antibodies could account for the remaining translation, that is inhibited by 2A-cp^N^. Alternatively, as eIF4G1 stabilizes eIF4E on m^7^GTP caps, it is possible that this cleavage product stabilizes eIF4E on caps and prevents any alternative modes of initiation, such as eIF3D-dependent translation. These possibilities are discussed below but await further validation.

**Figure 3.**
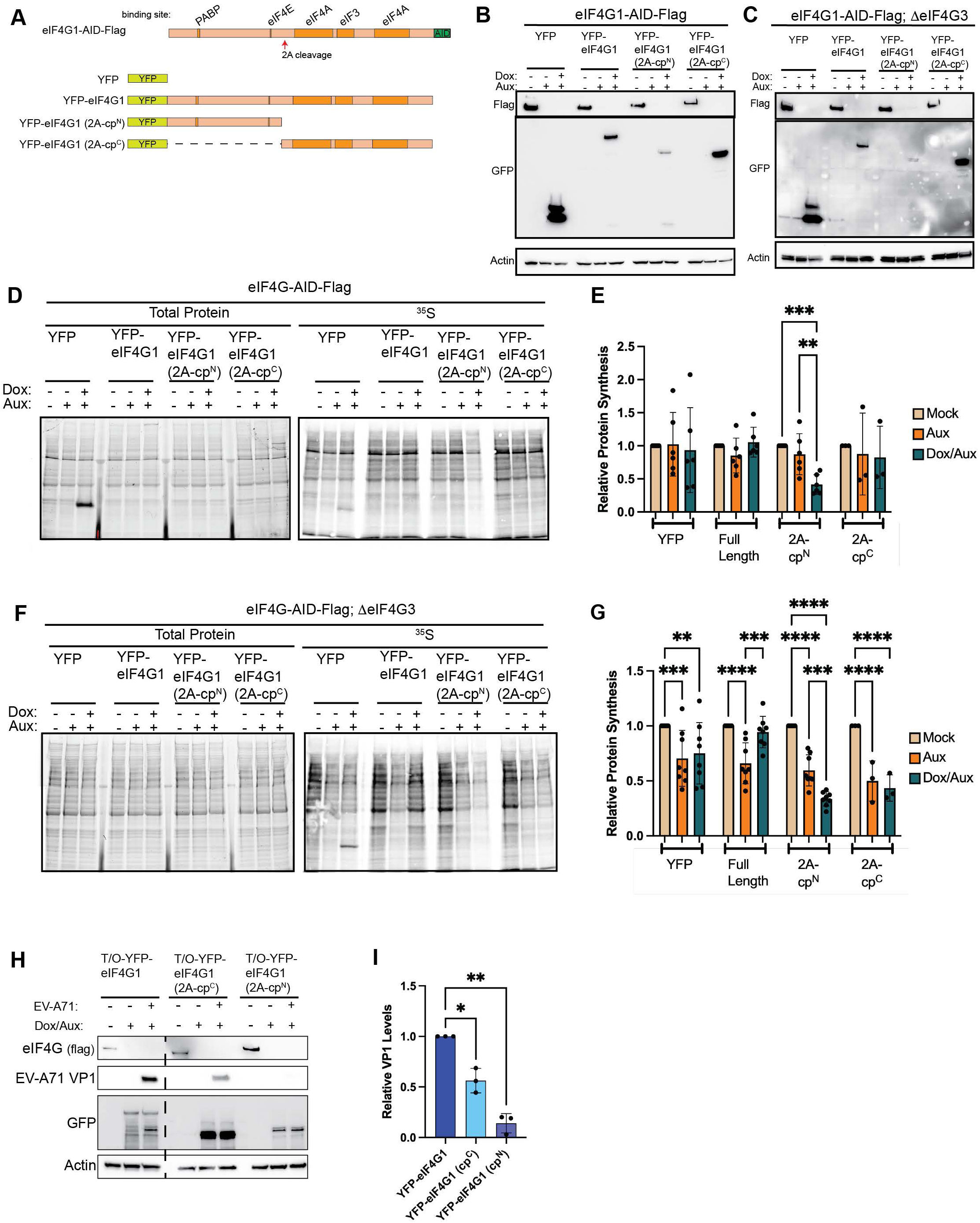
The N-terminal 2A protease cleavage fragment of eIF4G (2A-cp^N^) acts as a dominant translational repressor. (A) Schematic of tet-on YFP-eIF4G constructs designed to express 2A-mediated cleavage products of eIF4G. (B & C) Exogenous cleavage products or full-length eIF4G1 are induced upon addition of doxycycline in eIF4G1-AID-Flag (B) and eIF4G1-AID-Flag;∆eIF4G3 (C) cells. Co-treatment with auxin effectively depletes endogenous eIF4G1. (D & E) 2A-cp^N^ inhibits cellular protein synthesis in cells where eIF4G3 provides translational buffering, while 2A-cp^C^ has no effect. (F & G) 2A-cp^N^ inhibits cellular protein synthesis in the absence of eIF4G3, when translation is not buffered by this paralog; exogenous full-length eIF4G1 rescues translation in this context. (H & I) 2A-cp^C^ is not sufficient to fully rescue EV-A71 protein expression, while 2A-cp^N^ severely suppresses it, suggesting that 2A-cp^N^-mediated translational inhibition contributes to regulation of viral fitness. Statistical significance determined by two-way ANOVA (E & G) or one-way ANOVA (I) with Geisser-Greenhouse correction and Dunnett’s multiple comparisons test with individual variances. Mean ± SD, n≥3. *p<0.0332, **p<0.0021, ***p<0.0002, ****p<0.0001.

However, this does raise the possibility that 2A-cp^N^ has a role in regulating gene expression, particularly during enterovirus infection. It is well established that 2A-cp^C^ binds to IRES elements to recruit 43S PICs to viral mRNAs^19^, but the role of 2A-cp^N^ has not been explored. To test whether 2A-cp^N^ contributes to viral replication independently of 2A-cp^C^, we infected depleted and rescued cells with Enterovirus A 71 (EVA-71) and monitored viral protein production by western blotting for VP1 at 9 hours post-infection (**Fig. 3H & I**). As expected, full-length eIF4G1 supported robust viral protein production, while 2A-cp^N^ alone did not, consistent with its lack of the MIF4G domain required for IRES-mediated 43S recruitment. Notably, 2A-cp^C^ only partially rescued viral protein production, suggesting that the translational repressive activity of 2A-cp^N^ contributes to efficient viral replication beyond what IRES-driven translation alone can achieve. These results suggest that 2A-cp^N^ rewires cellular translation during enterovirus infection to promote viral replication.

Viral proteases are not unique in their ability to cleave eIF4G. The cellular cysteine-aspartic protease caspase-3 (casp3), activated during stress responses associated with apoptosis, also cleaves eIF4G at two sites distinct from those targeted by enteroviral 2A protease ^10,11^. Casp3-mediated cleavage generates three fragments, which we term casp3-cp^N^ (N-terminal), casp3-cp^M^ (middle), and casp3-cp^C^ (C-terminal) (**Figure 4A**). Based on known interaction domains, casp3-cp^N^ retains PABP-binding capacity, casp3-cp^M^ retains binding to eIF4E and contains the MIF4G domain that binds eIF3, and one eIF4A interaction site, and casp3-cp^C^ retains the second eIF4A-binding site.

**Figure 4.**
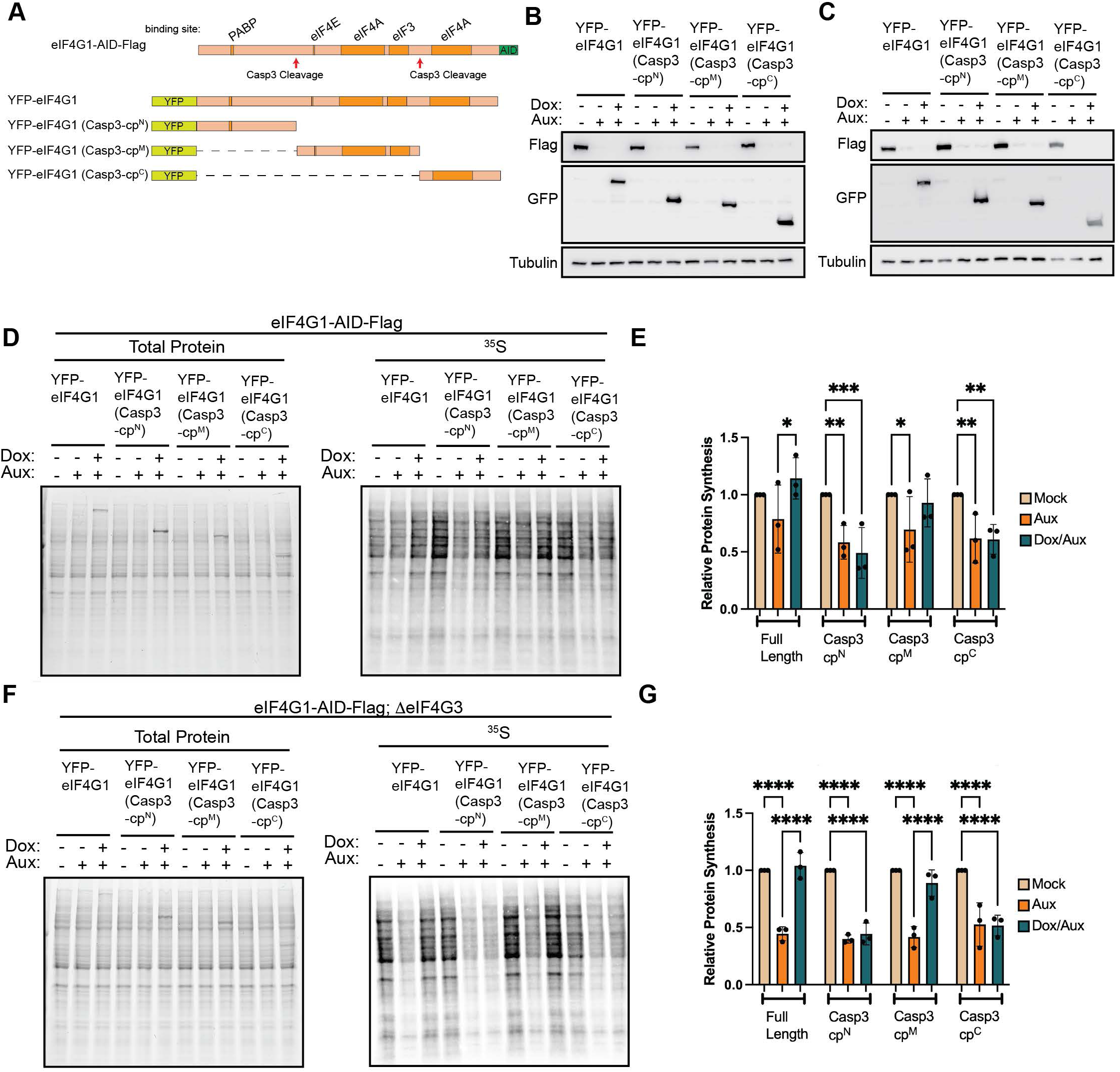
The middle caspase-3 cleavage fragment of eIF4G (casp3-cpM) fully supports basal translation. (A) Schematic of tet-on YFP-eIF4G constructs designed to express caspase-3 cleavage products of eIF4G. (B & C) Exogenous cleavage products or full-length eIF4G1 are induced upon addition of doxycycline in eIF4G1-AID-Flag (B) or eIF4G1-AID-Flag;∆1eIF4G3 (C) cells. Endogenous eIF4G1 is depleted upon co-treatment with auxin. (D–G) Global protein synthesis measured by [^35^S]-methionine incorporation following endogenous eIF4G1 depletion and rescue with the indicated exogenous constructs in eIF4G1-AID-Flag (D & E) or eIF4G1-AID-Flag;∆1eIF4G3 (F & G) cells. (D & E) Exogenous expression of full-length eIF4G1 or casp3-cp^M^ fully rescues translation in cells where eIF4G3 provides translational buLering. (F & G) Exogenous expression of full-length eIF4G1 or casp3-cp^M^ fully rescues translation in the absence of eIF4G3. Statistical significance determined by two-way ANOVA with Geisser-Greenhouse correction and Dunnett’s multiple comparisons test with individual variances (E & G). Mean ± SD, n≥3. *p<0.0332, **p<0.0021, ***p<0.0002, ****p<0.0001.

To determine whether casp3-generated fragments can regulate protein synthesis independently of caspase activation, we generated doxycycline-inducible casp3 cleavage products in eIF4G1-AID-Flag (**Figure 4B)** and eIF4G1-AID-Flag;∆eIF4G3 cells (**Figure 4C**). Using the same strategy as for 2A cleavage products, endogenous eIF4G1 was acutely depleted and replaced with individual casp3 cleavage products. In contrast to the inhibitory effect of the enteroviral 2A-cp^N^, none of the casp3-generated fragments repressed global protein synthesis (**Figure 4D – G**). Instead, expression of casp3-cp^M^, which retains binding to eIF4E, eIF3, and eIF4A, fully restored protein synthesis in eIF4G1-AID-Flag (**Figure 4D & E**) and eIF4G1-AID-Flag;∆eIF4G3 (**Figure 4F & G**).

Although these assays do not resolve effects on individual mRNAs, they indicate that PABP binding and the secondary eIF4A-binding domain are dispensable for sustaining bulk translation. These findings are consistent with prior *in vitro* studies demonstrating that the central domain of eIF4G constitutes the core translational scaffold ^51^ and RNA tethering studies monitoring the translation of reporter mRNAs in cells^52^. However, we believe this is the first demonstration that global translation can be sustained by this central fragment. Importantly, these findings challenge the prevailing model of translation initiation, which posits that formation of a closed-loop mRNP is required for efficient protein synthesis ^3,4^. Notably, casp3-cp^M^ lacks the PABP-binding domain and therefore cannot support closed-loop mRNP formation.

To more directly test this, we generated full-length eIF4G1 rescues with point mutations that ablated binding to PABP or eIF4E (**Figure 5A**). Previous work has established the motif necessary for PABP binding as ^174^KRERKIRIRD^184^ ^53^. Mutagenesis of this region ablates PABP binding by eIF4G1 ^54^. Similarly, it is established that the canonical eIF4E binding sequence is YXXXXLɸ where ɸ is any hydrophobic residue ^55,56^. Full interrogation of each amino acid has not been completed; although, far-western blot analysis suggested that Y612A mutation could fully ablate eIF4E interaction^56^. We generated full-length eIF4G1 that was previously demonstrated to be incapable of binding to PABP (eIF4G∆PABP) and two mutants that were prediction to disrupt eIF4E binding: a single Y612A mutant or full mutagenesis of every amino acid in the binding motif to alanine (eIF4G∆eIF4E) (**Figure 5A**). These mutations were also combined to generate exogenous eIF4G1 constructs with PABP and eIF4E binding mutations together. As eIF4G1-AID-Flag;∆eIF4G3 cells showed the most robust inhibition and rescue by full length eIF4G, we focused on this cell line. As before, co-treatment of auxin and doxycycline resulted in expression exogenous constructs (**Figure 5B**). We began by testing the efficacy of our mutations to ablate binding to individual proteins. As in previous studies, eIF4G1 constructs carrying the ∆PABP mutation (∆PABP, ∆PABP/∆eIF4E and ∆PABP/Y612A) fail to stably interact with PABP (**Figure 5C & D**). Similarly, constructs carrying the ∆eIF4E mutation fail to interact with eIF4E (∆eIF4E, ∆PABP/∆eIF4E) fully lose the ability to interact with eIF4E. However, we found that the Y612A mutation only partially reduced the ability of eIF4G1 to interact with eIF4E, resulting in a 50% reduction in stable interaction. Thus, this construct functions as a hypomorph for eIF4E binding.

**Figure 5.**
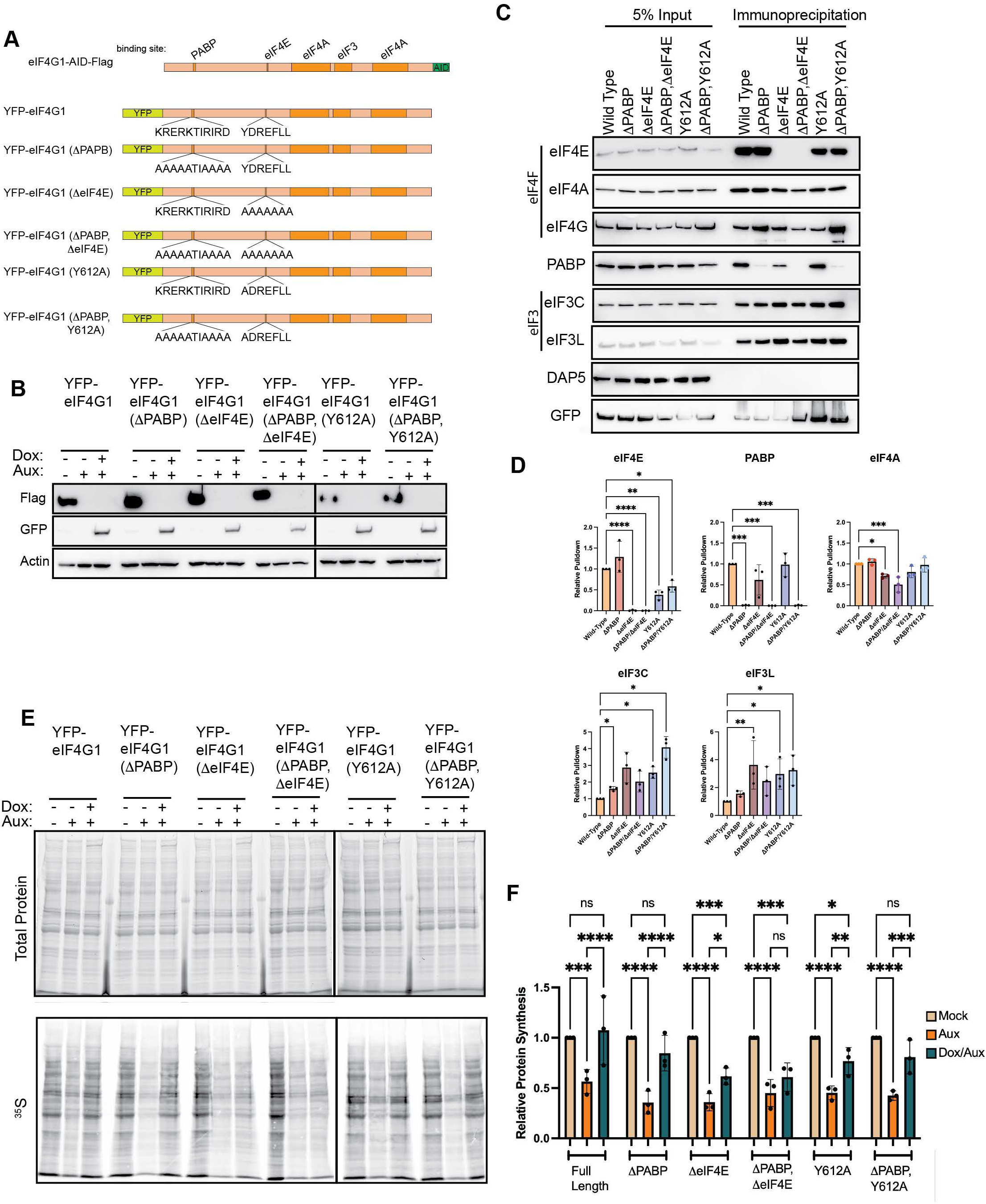
PABP interaction is dispensable for eIF4G-mediated translation. (A) Schematic of tet-inducible YFP-tagged wild-type and mutant eIF4G rescue constructs. ∆PABP and ∆eIF4E mutations replace the respective binding motifs with alanine substitutions (KRERKTIRIRD>AAAATIAAAA and YDREFLL>AAAAAAA, respectively); Y612A is a point mutation in the eIF4E binding site. (B) Western blotting confirms exogenous YFP-tagged eIF4G constructs are induced by doxycycline and endogenous eIF4G1-AID-Flag is simultaneously depleted by auxin in cells lacking eIF4G3-mediated translational buffering. (C & D) GFP co-immunoprecipitation of YFP-tagged eIF4G constructs. eIF4E- and PABP-binding mutants selectively fail to co-immunoprecipitate eIF4E or PABP, respectively. Y612A shows reduced, but not abolished, eIF4E binding. All mutants retain association with eIF4A and eIF3. (E & F) Global protein synthesis measured by [^35^S]-methionine incorporation following endogenous eIF4G1 depletion and rescue with the indicated exogenous constructs. Full-length eIF4G and PABP-binding-deficient eIF4G (∆PABP) fully restore protein synthesis. In contrast, eIF4E-binding mutants (∆eIF4E; ∆PABP,∆eIF4E) fail to rescue, and the reduced-affinity Y612A mutant shows intermediate recovery, demonstrating that eIF4E, but not PABP, binding is required for eIF4G-mediated translational activity. Statistical significance determined by one-way ANOVA (D) or two-way ANOVA (F) with Geisser-Greenhouse correction and Dunnett’s Multiple comparisons test with individual variances. Mean ± SD, n≥3. *p<0.0332, **p<0.0021, ***<0.0002, ****p<0.0001.

We next tested these constructs’ ability to rescue translation upon eIF4G1 depletion (**Figure 5E & F**). As would be suggested by casp3 proteolytic fragments, loss of PABP binding (∆PABP) has no effect on the ability of exogenous eIF4G1 to rescue global translation. Loss of eIF4E interaction (∆eIF4E) renders exogenous construct incapable of rescuing translation; although, unexpectedly, the ∆eIF4E construct provided a statistically significant partial rescue of global translation, and the ∆PABP,∆eIF4E double mutant trended in the same direction, though this did not reach statistical significance. These results suggest that eIF4G1 retains residual translational activity independent of eIF4E engagement. One possible explanation is that eIF4G1 is recruited to a subset of mRNAs independently of eIF4E through the RNA-binding activity of its HEAT1 domain. We and others have previously demonstrated that eIF4G1 HEAT1 domain possesses intrinsic RNA binding activity^57–61^, including affinity for G-quadruplex structures, raising the possibility that direct RNA recruitment of eIF4G1 contributes to cap-independent translation of specific transcripts. Finally, we found that Y612A mutations fully rescued translation despite the reduction in binding suggesting that a 50% reduction in interaction is still sufficient to drive the majority of cellular translation.

Given these data, we next sought to determine if 2A-cp^N^’s ability to inhibit translation was dependent upon its ability to interact with PABP, eIF4E, or both. We generated inducible 2A-cp^N^ constructs with the inability to bind PAPB (2A-cp^N^, ∆PABP), eIF4E (2A-cp^N^, ∆eIF4E) or both (2A-cp^N^, ∆∆eIF4E/PABP) (**Figure 6A**). These constructs were expressed in eIF4G1-AID-Flag and eIF4G1-AID-Flag;∆eIF4G3 cells (**Figure 6B & C**). As before, we monitored protein synthesis after depletion or depletion and rescue using [^35^S] metabolic labeling (**Figure 6D - G**). Surprisingly, while PABP binding is dispensable rescue of basal translation (**Figure 4, 5**), we found that 2A-cp^N^ must be able to bind PABP to effectively inhibit translation. As expected, failure to interact with eIF4E also ablates the inhibitory activity of 2A-cp^N^ in the presence (**Figure 6D & E**) or the absence (**Figure 6F & G**) of eIF4G3. These results demonstrate that the inhibitory complex requires closed-loop formation, while active translation can dispense of this mRNP conformation.

**Figure 6.**
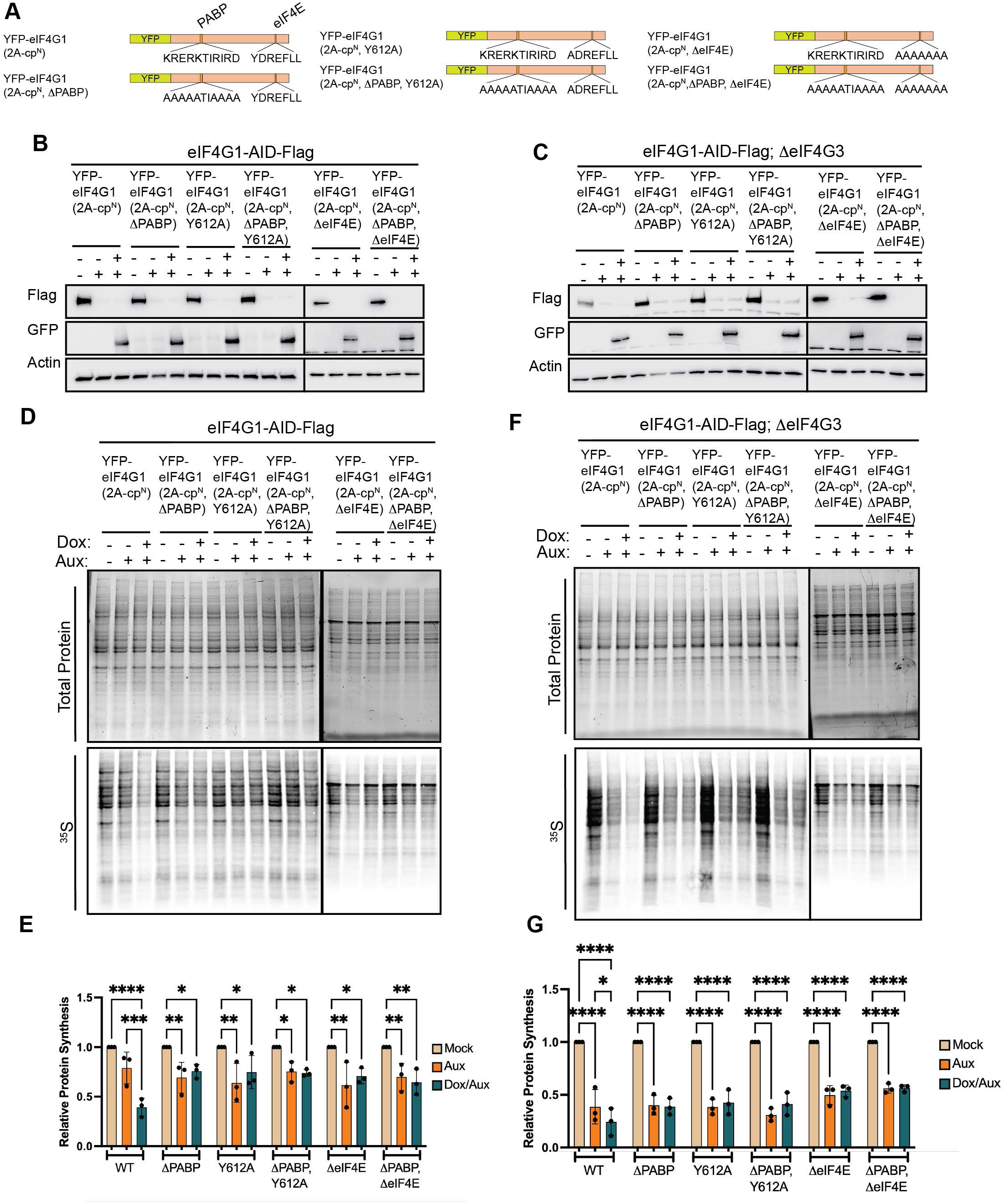
PABP binding is required for 2A-cp^N^-mediated translation repression. (A) Schematic of exogenous tet-on mutant 2A-cp^N^ constructs. (B & C) Exogenous expression of mutant 2A-cp^N^ constructs upon addition of doxycyline and depletion of endogenous eIF4G upon addition of auxin in eIF4G1-AID-Flag (B) and eIF4G1-AID-Flag;∆eIF4G3 (C) cells. (D – G) Global protein synthesis measured by [^35^S]-methionine incorporation following endogenous eIF4G1 depletion and rescue with the indicated exogenous constructs in eIF4G1-AID-Flag (D & E) or eIF4G1-AID-Flag;∆eIF4G3 (F & G) cells. In contrast to stimulatory activity, for which PABP binding is dispensable, inhibitory activity by 2A-cp^N^ requires PABP interaction in both genetic backgrounds. Statistical significance determined by two-way ANOVA with Geisser-Greenhouse correction and Dunnett’s Multiple comparisons test with individual variances (E & G). Mean ± SD, n≥3. *p<0.0332, **p<0.0021, ***<0.0002, ****p<0.0001.

These lack of closed-loop mRNP formation to effect robust translation has been suggested by other data. In yeast, loss of Pab1, the yeast ortholog of human PABP, primarily reduces mRNA stability without affecting translational efficiency ^62^. Humans have multiple paralogs of PABP, with most critical being PABPC1 and PABPC4 (**Supplemental Figure 3A**). Using a DHFR degron, previous work has shown that, similar to yeast, the primary effect of loss of both proteins is destabilization of mRNAs^63^. Using the DHFR system, full depletion of PABPC1 was possible at 8 hours and most experiments using this system were conducted beyond this time point. We adapted our AID approach as it allows for more rapid decay kinetics to better delineate immediate translation effects from later mRNA stability effects. To this end, we knocked out PABPC4 and tagged PABPC1 with the same AID tag except we added a EGFP tag, rather a flag tag (**Figure 7A - B**). As with eIF4G1-AID-Flag cells, neither PABPC1-AID-EGFP, nor PABPC1-AID-EGFP;∆PABPC4 cells have any defect regarding basal protein synthesis (**Figure 7C & D**). Upon addition of auxin, we found significantly improved decay kinetics relative to the DHFR system with nearly full depletion achieved by 2 hours post-auxin treatment as monitored by western blotting to PABP or GFP in both PABPC1-AID-EGFP and PABPC1-AID-EGFP;∆PABPC4 cells (**Figure 7E & F**). Concurrent with depletion of PABPC1, we found a time-dependent increase of PABPC4 in our PABPC1-AID-EGFP lines. Previous studies have shown that PABPC1 mRNA is autoregulated by an adenosine-rich sequence in its 5’UTR ^64,65^. High levels of PABP protein repress PABP mRNA translation, which is relieved as PABP levels decrease. Our analysis of PABPC4 mRNA 5’UTR reveals a similar adenosine-rich sequence which likely regulates synthesis in a similar manner (**Supplemental Figure 3B**). Thus, upon auxin mediated depletion of PABPC1, the synthesis of PABPC4 is likely stimulated to compensate, suggesting a larger network of regulation between PABP paralogs; however, this confirmation of this hypothesis awaits further testing.

**Figure 7.**
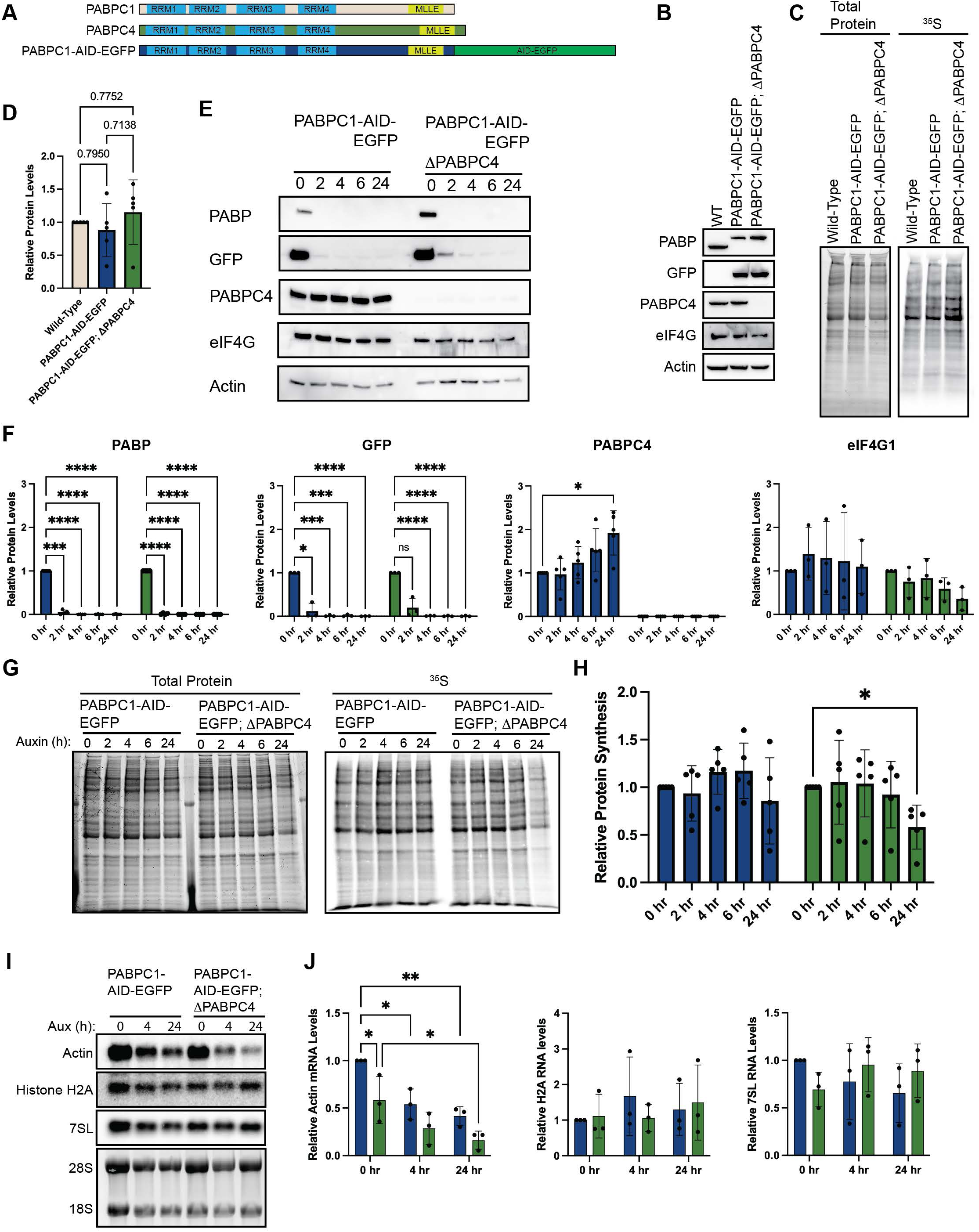
PABP is not required for bulk protein synthesis but is required for mRNA stability. (A) Schematic of PABPC1, PABPC4, and AID-EGFP-tagged PABPC1, indicating RNA recognition motifs (RRM1–4), MLLE domain, and position of the AID-EGFP insertion. (B) Western blot validation of cell lines used in this figure. CRISPR-mediated AID-EGFP tagging of PABPC1 is confirmed by mobility shift and anti-GFP immunoreactivity; CRISPR-mediated PABPC4 knockout is confirmed by loss of anti-PABPC4 signal. (C & D) Basal global protein synthesis measured by [^35^S]-methionine incorporation in wild-type, PABPC1-AID-EGFP, and PABPC1-AID-EGFP;∆PABPC4 cells. Neither AID-EGFP tagging of PABPC1 nor PABPC4 knockout alters protein synthesis under steady-state conditions. (E & F) Auxin time-course (0, 2, 4, 6, 24 hours) in PABPC1-AID-EGFP and PABPC1-AID-EGFP;∆PABPC4 cells. Anti-PABP and anti-GFP blots confirm rapid depletion of PABPC1-AID-EGFP within 2 hours in both backgrounds. PABPC1 depletion induces a compensatory increase in PABPC4 protein in cells retaining PABPC4, consistent with autoregulatory buffering. eIF4G levels are unaffected in either background. (G & H) Global protein synthesis measured by [^35^S]-methionine incorporation across the same auxin time-course. Neither acute (2–6 hours) nor chronic (24 hours) loss of PABPC1 alone affects protein synthesis. Acute co-depletion of PABPC1 and PABPC4 similarly has no effect; however, chronic co-depletion results in a significant reduction in protein synthesis, consistent with downstream loss of mRNA template rather than a direct effect on translational efficiency. (I & J) Northern blot analysis of polyadenylated actin mRNA, non-polyadenylated histone H2A mRNA, and non-polyadenylated 7SL ncRNA following acute (4 hours) or chronic (24 hours) auxin treatment; 28S and 18S rRNA serve as loading controls. Actin mRNA levels decline progressively upon PABPC1 depletion, with significantly greater reduction in the PABPC1-AID-EGFP;∆PABPC4 background, indicating that PABPC4 compensates for PABPC1 loss to maintain polyadenylated mRNA stability. Non-polyadenylated transcripts are unaffected by loss of either PABPC1 or PABPC4. Statistical significance determined by one-way ANOVA (D) or two-way ANOVA (F, H, J) with Geisser-Greenhouse correction and Dunnett’s Multiple comparisons test with individual variances. Mean ± SD, n≥3. *p<0.0332, **p<0.0021, ***<0.0002, ****p<0.0001.

We find no alterations to basal protein synthesis upon PABPC1 depletion in either PABPC1-AID-EGFP nor PABPC1-AID-EGFP;∆PABPC4 cells through 6 hours of depletion (**Figure 7G & H**). However, chronic loss (24 hours) of PABPC1 in cells lacking PABPC4 results in a ∼50% reduction in protein synthesis. These data are consistent with previous work that suggests that PABP primarily regulates protein synthesis by promoting mRNA stability rather than stimulating translation. Additionally, baseline levels of PABPC4 and increased levels upon PABPC1 depletion are sufficient to retain mRNA stability. To confirm this, we conducted northern blotting on the polyadenylated Actin mRNA and the non-polyadenylated Histone H2A mRNA following PABPC1 depletion in the presence or absence of PABPC4. We found that the actin mRNA had reduced levels at steady state in ∆PABPC4 cells, indicating that it plays a normal role in mRNA stability. Depletion of PABPC1 rapidly reduced the steady state levels of the Actin mRNA. This effect was significantly enhanced in the absence of PABPC4 (**Fig 7I & J**). In contrast, the non-polyadenylated histone H2A mRNA was unaffected by knockout of PABPC4 or depletion of PABPC1 in either cellular context. Collectively, these results strongly suggest that the reduction in protein synthesis observed in PABPC1-AID-EGFP;∆PABPC4 cells at 24 hours post depletion is due to reduced levels of cellular mRNAs, rather than acute effects on translation initiation.

## Discussion

The closed-loop model of translation posits that eIF4E-eIF4G-PABP bridging between the 5′ cap and the 3′ poly(A) tail is a central driver of efficient protein synthesis ^3,4^. This model has been influential in framing our understanding of cap-dependent initiation for nearly three decades. Here, we present data that fundamentally challenge it. Using acute, AID-mediated depletion of eIF4G1 combined with systematic replacement by defined cleavage products and interaction mutants, we demonstrate that PABP binding by eIF4G is dispensable for sustaining global translation. We further show that the closed-loop architecture, while not required for productive initiation, can be co-opted by enteroviral proteolytic fragments to actively repress host protein synthesis. As detailed in **Figure 8**, these findings refine the canonical view: eIF4G-PABP bridging is dispensable for bulk translation initiation but remains a functionally important interface that that may modulates mRNA-specific translation efficiency and, under conditions of viral infection, is exploited for translational silencing.

**Figure 8.**
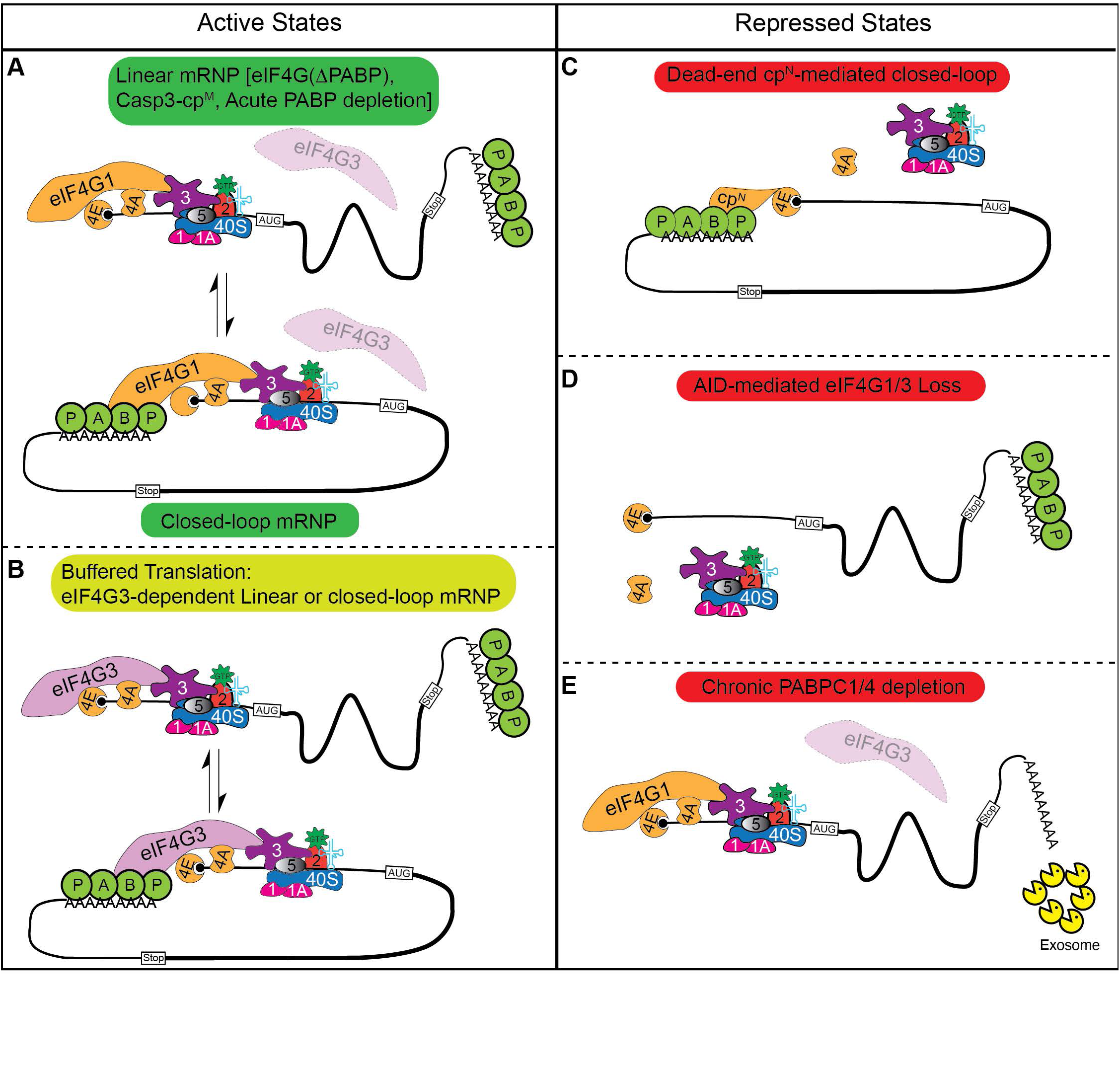
Topologies of eIF4G-dependent translation. (A & B) Active translation is supported by eIF4G1 (A) or, when eIF4G1 is limiting, by eIF4G3 (B). Prior studies established that active mRNPs can adopt a closed-loop conformation in which eIF4G bridges eIF4E at the 5′ cap and PABP at the poly(A) tail. Our data demonstrate that productive translation also occurs on linear mRNPs when eIF4G is unable to interact with PABP or when PABP is acutely depleted with 43S PIC recruitment mediated by the MIF4G domains of eIF4G1 or eIF4G3. (C - E) Repressive states arise through three distinct mechanisms: formation of a dead-end closed-loop mRNP by 2A-cp^N^, which bridges eIF4E and PABP but lacks a functional MIF4G domain and cannot recruit 43S PICs (D) ; complete loss of both eIF4G1 and eIF4G3, eliminating the scaffold for 43S PIC recruitment (E) ; or chronic depletion of PABP proteins (E). The reduction in protein synthesis observed under chronic PABP depletion reflects decreased mRNA abundance rather than a direct impairment of translational efficiency.

### Closed-loop mRNP formation is not required for global protein synthesis

Several independent lines of evidence support this reinterpretation. First, the middle caspase-3 cleavage fragment of eIF4G (casp3-cp^M^), which retains the eIF4E, eIF3, and eIF4A binding sites but entirely lacks the PABP-interaction domain, is sufficient to fully rescue global protein synthesis upon acute eIF4G1 depletion in agreement with previous *in vitro* biochemical studies and mRNA tethering assays (**Figure 4)** ^51,52,66^. Second, full-length eIF4G1 carrying a PABP-binding mutation (∆PABP) rescues translation equivalently to wild-type eIF4G1, demonstrating that even in the context of intact eIF4G1, PABP engagement is not required to support bulk protein synthesis (**Figure 5**). Third, acute depletion of both PABPC1 and PABPC4, the major cytoplasmic PABP paralogs, produces only a modest and delayed reduction in global translation, with the primary consequence being mRNA destabilization rather than an acute initiation defect (**Figure 7**). Taken together, these results indicate that eIF4E-eIF4G1-PABP bridging is not a prerequisite for productive cap-dependent translation initiation in mammalian cells. The modest effect of PABP depletion on bulk translation is consistent with the view that the poly(A) tail confers a competitive advantage among mRNAs in a crowded cellular environment, as suggested by the observation that poly(A) tail effects are more pronounced *in vivo* than in cell free systems, rather than constituting a universal requirement for initiation. Whether the closed-loop modulates relative translation efficiency among specific mRNA subsets remains an important open question that ribosome profiling will be needed to resolve. It is worth noting that our data only report on bulk global protein synthesis. It is conceivable that there are specific mRNAs that have an outsized need for closed-loop mRNP to sustain productive synthesis. Our data would suggest that such mRNAs are the exception rather than the rule, though this prediction awaits direct testing.

In contrast, we find that this same closed-loop architecture is actively harnessed to silence translation. The N-terminal eIF4G cleavage product generated by enteroviral 2A protease (2A-cp^N^) is a potent, dominant translational repressor, and its inhibitory activity requires both its eIF4E-binding and PABP-binding domains. Disrupting either interaction is sufficient to abolish repression. This indicates that 2A-cp^N^ functions by simultaneously engaging the cap and poly(A) tail to form a closed-loop mRNP that prevents productive 43S PIC recruitment by sequestering eIF4E and PABP without providing the eIF3 and eIF4A binding activities needed for ribosome loading. The result is an mRNA that is topologically closed but translationally inert. In this respect, 2A-cp^N^ likely functions similarly to the eIF4E-binding proteins (4EBPs), which sequester eIF4E to block eIF4G association and prevent initiation complex assembly. This co-option of the closed-loop architecture as a repressive mechanism provides a resolution to an outstanding paradox: namely, why enteroviral infection produces near-complete translational shutoff while simple depletion of eIF4G does not. Our data suggest that enterovirus cleavage results not only in the loss of eIF4G scaffolding activity but in the gain of active repressive function by 2A-cp^N^.

These findings also invite a reexamination of the endogenous function of eIF4G-PABP interaction. Rather than dismissing closed-loop mRNP formation as functionally irrelevant, our data suggest that its physiological roles are more closely tied to post-initiation steps than to driving ribosome recruitment. Indeed, two well-established functions of eIF4G-PABP bridging lie outside of initiation. First, the eIF4G-PABP interaction has been shown to suppress nonsense-mediated mRNA decay (NMD)^54^, coupling the translation machinery to mRNA surveillance and quality control. Second, recent biochemical work using a reconstituted mammalian translation system demonstrated that eIF4F potently stimulates translation termination by enhancing eRF3 GTPase activity and facilitating release factor dissociation following peptide release^67^. In that study, the PABP-eRF3 interaction was proposed to recruit eIF4F to the terminating ribosome via closed-loop formation, implicating this architecture in coordinating the handoff between termination and ribosome recycling. Importantly, this function was shown to be dependent upon the central domain of eIF4G that is analogous to Casp3-cp^M^, which we show is sufficient to rescue bulk translation.

Together, these observations suggest that eIF4G-PABP bridging primarily functions at post-initiation stages. This interaction may be more relevant in promoting termination fidelity and suppressing aberrant mRNA decay rather than serving as a direct driver of ribosome recruitment. The translational stimulation historically attributed to poly(A)-tail-dependent closed-loop formation may largely reflect PABP’s role in protecting mRNAs from premature degradation and maintaining the pool of translatable mRNA, rather than a direct kinetic enhancement of initiation.

### eIF4G3 buffers basal translation and can be activated upon eIF4G1 loss

A second major finding of this study is that basal translation is substantially buffered against acute eIF4G1 loss by its paralog eIF4G3, which can sustain a portion of global protein synthesis under homeostatic conditions and is further activated when eIF4G1 is compromised. This provides a compelling explanation for the modest translational defects observed in prior siRNA-based studies of eIF4G1 knockdown ^48,68^: the extended timescales required for RNAi likely allowed eIF4G3 to compensate before measurements were taken. This highlights that AID-mediated depletion can reveal previously unappreciated aspects of biology. By achieving depletion within hours, we can separate the initial eIF4G3-independent translational response from eIF4G3-mediated buffering and show that both components are real and distinct. The fact that enteroviruses evolved to cleave eIF4G3 following eIF4G1, and that this second cleavage correlates with more complete translational shutoff ^42,69^, provides independent biological support for eIF4G3’s buffering role

Notably, this buffering occurs without detectable upregulation of eIF4G3 protein levels, implying that a pool of eIF4G3 exists in a less active state under normal conditions and can be mobilized when eIF4G1 is limiting. This contrasts with our demonstration that PABPC4 likely compensates for loss of PABPC1 through increasing its abundance. How eIF4G3 activity is normally restrained remains an open question. Competition with eIF4G1 for shared binding partners (e.g., eIF4E, eIF4A, eIF3) or differential post-translational regulation could each contribute. Notably, the eIF4E-binding domain is 100% conserved between eIF4G1 and eIF4G, arguing against differential affinity for eIF4E1 as the mechanistic basis for preferential eIF4G1 activity under basal conditions. If eIF4G3 is regulated through its eIF4E interaction, that regulation must be conferred through auxiliary binding domains rather than the YXXXXLɸ motif. Biophysical data has revealed additional eIF4E binding motifs within eIF4G downstream of the canonical binding site ^70^, but no work has been conducted to determine if they play similar roles in eIF4G3; however, the amino acids required for binding in eIF4G (SDVVLD) are conserved in eIF4G3. The differential sensitivity of eIF4G1 and eIF4G3 to viral proteases ^42^ and cellular caspases ^46^ suggests that paralog-selective regulation is a feature of physiological translational control and that conditions which specifically impair eIF4G1 may unmask latent eIF4G3 activity. Consistent with a broadly supportive but redundant role, constitutive loss of eIF4G3 alone does not affect basal protein synthesis (**Figure 2**), establishing that eIF4G1 is sufficient in the absence of its paralog. This functional architecture in which eIF4G1 is dominant but eIF4G3 can compensate under stress may represent a general organizational principle of the initiation apparatus that provides robustness against acute perturbations. Indeed, previous data has demonstrated that eIF4G3 can incorporate into non-canonical eIF4F complexes under hypoxic stress through interaction with eIF4E2, a paralog of eIF4E ^26,50^. Given the 3 major paralogs of eIF4E (eIF4E, eIF4E2, eIF4E3) and the two eIF4E-interacting eIF4G paralogs (eIF4G, eIF4G3) it is conceivable that 6 distinct eIF4F complexes could form. How the formation of these would be regulated and whether each supports the translation of distinct subsets of mRNAs awaits further research; although, our data suggests that activation of eIF4G3 is not restricted solely to hypoxic conditions.

Our data also confirm previous work ^63^ demonstrating that PABPC4 is a functional paralog of PABPC1. Upon loss of PABPC1, PABPC4 can compensate by stabilizing mRNAs (**Figure 7**). In contrast to eIF4G3, which can compensate and buffer translation independent of an increase in protein levels, PABPC4 levels are elevated in response to PABPC1 depletion. This is likely achieved through autoregulatory adenosine-rich motifs in the 5’UTR of both mRNAs. Analysis of the 5’UTRs of other PABP paralogs reveal that they lack such motifs. This is likely due to their specialize roles in development or their tissue restricted nature. These two systems thus represent distinct buffering strategies. One relies on activating pre-existing, but latent protein pools (eIF4G3), while the other relies on a functional increase in protein levels (PABPC4). It will be interesting to determine how broadly each strategy is deployed across the initiation machinery, particularly given that eIF4E and eIF4A possess several paralogs each.

### Enteroviral eIF4G cleavage products have diverse functions

Our results suggest a separation of function between the two fragments generated by enteroviral 2A protease that has not previously been appreciated in full. While it has long been established that 2A-cp^C^ supports viral IRES-driven translation by engaging eIF3 and eIF4A^16–18^, the functional significance of 2A-cp^N^ has been largely unexplored. We now show that 2A-cp^N^ actively suppresses cap-dependent host translation through closed-loop mRNP sequestration. Together, these activities constitute a two-pronged strategy in which a single proteolytic cleavage event simultaneously redirects the translational machinery toward viral mRNAs and silences competing host transcripts with each fragment contributing a distinct activity. Consistent with this model, eIF4G cleavage has been associated with enrichment of viral mRNA in polysomes^71^, and our data suggest that 2A-cp^N^-mediated repression of host translation is an active contributor to this redistribution. Indeed, expression of 2A-cp^C^ in the absence of 2A-cp^N^ does not fully rescue infection by EV-A71. These data suggest that 2A-cp^N^ plays an active role in rewiring cellular translation during infection. It is intriguing that further blocking cap-dependent translation after cleavage of eIF4G can exert this effect. It is possible 2A-cp^N^ can stabilize eIF4E on m^7^GTP caps as eIF4E’s residence on m7GTP caps is stabilized by eIF4G interaction^72^. In fact, pioneering biophysical studies in yeast demonstrated this phenomenon used fragments of eIF4G rather than full length protein^73,74^. Thus, 2A-cp^N^ could stably generate a dead-end product that prevents 43S recruitment by other means. eIF3D can also bind the m^7^GTP cap and competes with eIF4E for access to this structure^75^. During persistent cellular stress, eIF3d is activated through a phosphorylation switch to drive translation of specific pro-survival mRNA subsets, including regulators of the integrated stress response^76^. It is conceivable that 2A-cpN stabilizes eIF4E on these mRNAs and physically occludes cap access by eIF3d, preventing handoff to this alternative mode of initiation. However, future studies are needed to determine the interactions between 2A-cp^N^, eIF3D and viral infection.

### Caspase-3 cleavage of eIF4G may rewire rather than simply silence translation

In contrast to enteroviral proteolysis, caspase-3-mediated eIF4G cleavage generates fragments that are not repressive. The middle fragment fully rescues global translation, and neither the N-nor C-terminal fragments suppress protein synthesis (**Figure 4**). The capacity of casp3-cp^M^ to sustain translation invites a reconsideration of what caspase-3-mediated eIF4G cleavage actually accomplishes during cellular stress. Caspase-3 activation can have opposing outcomes in cells depending upon the magnitude of activation ^13^. Robust caspase-3 activation drives apoptosis, but low levels of caspase-3 activation can actually promote survival^7,8^.

Under such conditions, casp3-cp^M^, which retains eIF4E and eIF4A interactions but lacks the PABP-binding domain required for closed-loop mRNP formation, could sustain translation. Thus, caspase-3-mediated cleavage may remodel rather than silence the translational machinery by sustaining protein synthesis in a closed-loop-independent configuration during stress. As suggested by previous work, this could reduce the efficiency of translation termination ^67^ or enhance the susceptibility to NMD^54^. Moreover, while bulk protein synthesis is rescued by casp3-cp^M^ and eIF4G(∆PABP), it is likely that there are specific transcripts that have a higher dependence on eIF4G-PABP interaction, which may be specifically altered upon caspase-3 cleavage. This is in striking contrast to enteroviral 2A cleavage, where the equivalent N-terminal fragment actively sequesters eIF4E and represses host translation. Thus, eIF4G1 could be co-opted by viral and cellular proteases for opposite translational outcomes, with cleavage site choice determining functional consequence.

## Conclusions

This study provides mechanistic insight into translational regulation at multiple levels. We show that eIF4G3 acts as a physiological buffer that sustains global protein synthesis when eIF4G1 is impaired, that the central domain of eIF4G is sufficient for bulk translation without PABP interaction, and that enteroviral proteolysis generates a dominant repressor (2A-cp^N^) that co-opts closed-loop mRNP formation to silence host translation. These findings refine our understanding of eIF4G-PABP interactions: rather than serving as a central driver of productive initiation, this interface appears to function primarily at post-initiation stages, such as mRNA quality control, termination and stability, while also being exploitable for translation repression by viral proteolytic fragments. The closed-loop architecture is neither universally stimulatory nor universally repressive; its functional consequences depend on which proteins form it and what activities they carry. More broadly, the functional divergence between viral and cellular eIF4G cleavage products illustrates how the modular architecture of eIF4G enables opposing translational outcomes to be achieved through selective proteolytic processing.

## METHODS

### Cell Culture and Drug Treatments

HeLa or A549 cells were maintained in DMEM (Gibco) supplemented with 10% fetal bovine serum (Atlas Biologicals) and 1% Penicillin/Streptomycin (Corning) in a 37°C incubator at 5% CO_2_. Where indicated, cells were treated with 100 μg/ml Auxin (3-Indolacetic acid; Millipore-sigma) or 10 μg/ml Doxycycline (Fisher).

### CRISPR-mediated editing of cells

pSH-EFIRES-P-AtAFB2-mCherry-Weak-NLS (Addgene #129717) or pSH-EFIRES-Z-AtAFB2-Myc were co-transfected with pX300-AAVS1 (Addgene #227272) into HeLa or A549 cells, respectively, to generate parental cell lines. HeLa cells were selected with puromycin and A549 cells were selected with Zeocin and single cell colonies were isolated by limiting dilution. Homology directed repair plasmids were generated by flanking a cassette of IAA7-Flag-IRES-HygR for eIF4G or IAA7-EGFP-IRES-NeoR for PABPC1 by approximately 250 bp of genome DNA flanking the stop codons. Guide RNAs targeting the 3’ end of the open reading frame of each gene were cloned into pCas-Guide (Origene) according to manufacturer’s instructions. HeLa or A549 cells were plated in a 6-well dish and the following day transfected with equal amounts of HDR plasmid and pCas plasmids. Cells were selected using hygromycin (eIF4G) or G418 (PABPC1). For knockouts, guide RNAs were designed using Atum software and cloned into lentiCRISPR-v2-Blast (Addgene #83480). Lentiviral particles were generated in 293T cells using psPAX3 and pMD.2 support plasmids. Target cells were transduced in antibiotic free media and selected the following day with blasticidin. Clones were isolated by limiting dilution and screened by western blotting followed by genotyping.

### Metabolic Labeling and Western Blotting

Cells were briefly starved for 10 minutes in methionine/cysteine free DMEM containing 10% FBS. Cells were labeled with 5 μCi EasyTag EXPRESS^35^S Protein Labeling Mix (Revvity) for 10 minutes. Cells were lysed in NP-40 Lysis buffer (20 mM Tris [pH 8.0], 150 mM NaCl, 0.5% NP-40) and cleared by centrifugation (5 min, 14 000 *x g*). Lysates were quantified by bradford colorimetric assay and equal amounts of proteins were loaded onto BioRad Stain-free SDS-PAGE gels. Total protein loading was determined using BioRad Stain-free technology. Proteins were transferred to nitrocellulose membranes, dried, and exposed to phosphor screen overnight before detection using GE typhoon. Stain-free gels and autoradiography were quantified using Bio-Rad Image lab. Membranes were rehydrated in TBST and subjected to western blotting. Chemiluminescent signal was detected using Bio-Rad ChemiDoc plus and analyzed using Bio-Rad ImageLab software. Western blotting signal was normalized to total protein.

### Northern Blotting

Northern blotting probes were generated using DNA oligonucleotides synthesized by Integrated DNA Technologies and resuspended in diH_2_O to a final concentration of 6 μM. To end label the DNA, 1 μL of 6 μM DNA oligonucleotide, 14 μL diH_2_O, 2 μL 10X T4 PNK buffer, 1 μL T4 PNK (NEB), and 5 μL of [γ-^32^P] ATP (3000 Ci/ml) (Revvity) were incubated at 37°C for 1 hr. Post incubation, 80 μL of diH_2_O was added to the reaction mix and unincorporated radioactive nucleotides were filtered from the final probe pool by gel filtration in a G-25 spin column (Cytiva) according to manufacturer’s protocols.

Cells were lysed using 0.8 M Guanidinium thiocynate, 0.4 M Ammonium thiocynate, 0.1 M Sodium Acetate [pH 5.0], 5% glycerol, 48% Phenol. RNA was extracted by addition of 0.5 volumes of 100% chloroform followed by centrifugation and precipitation of the aqueous phase with 2 volumes of 100% ethanol. 5-10 ug of RNA was run on a 1.2% Agarose gel, with 1X H-E buffer (20 mM HEPES, 1 mM EDTA [pH 7.8]) and 7% formaldehyde, in 1X H-E buffer at room temperature for 18 hrs at 55 V with free buffer exchange. The RNA was transferred to Amersham Hybond - N+ membrane (Cytiva) via passive evaporation and UV crosslinked at 260 nm with 1200 μJ (X100) twice in a Stratalinker. The membranes were then stained with methylene blue and prehybridized for 1 hr at 65°C with 10 mL of ULTRAhyb Ultrasensitive Hybridization Buffer (Invitrogen). The hybridization buffer was then replaced with 10 mL of fresh buffer and allowed to reach temperature of 65°C. Pre-prepared ^32^P end labeled radioactive probe in 10 μL aliquots were denatured at 95°C for 10 mins. 1 mL of pre-warmed hybridization buffer was added to the 10 μL of denatured probe and then added back to the hybridization tube. The probe was incubated for 1 hour at 65°C then the temperature was reduced to 37°C overnight. Blots were then washed with 2X SSC / 0.1% SDS two times for 15 mins at 42°C, wrapped in cling wrap, and exposed on a Molecular Dynamics Kodak Storage Phosphor Screen S0230 overnight. Storage screens were then imaged on a 2020 Amersham Typhoon Phosphor Imager (GE), images were processed and quantified with ImageLab and compiled with Adobe Illustrator 2023. Imaged blots were stripped and prepared for re-probing by pouring around 100 mL of boiling 0.1X SSC / 0.05% SDS and rocking on a shaker until reaching room temperature three times.

### Enteroviral Infection Studies

eIF4G1-AID-Flag;∆eIF4G cells harboring Tet-On-EYFP-eIF4G(WT), Tet-On-EYFP-eIF4G(cp^N^) or Tet-On-EYFP-eIF4G(cp^C^) were seeded at 2.5 x 10^5^ cells/well in 12-well plates and incubated overnight at 37°C containing 5% CO_2_. The next day, and 16 hrs prior to the start of viral infection, cell culture media was replaced with either 5% FBS-containing DMEM or with 5% FBS DMEM supplemented with auxin, or 5% FBS DMEM supplemented with both auxin and doxycycline. EV-A71 stocks were diluted in Opti-Mem (Gibco) to an MOI of 3 and applied to the HeLa cell cultures. Viruses were allowed 1hr to attach at 37°C and 5% CO_2_ before being removed and residual inoculum was removed by a series of PBS washes. 5% FBS DMEM containing the same additives as used during the initial pretreatment period were resupplied to the HeLa cell cultures and allowed to propagate at 37°C until the indicated time. Here, the cell culture media was removed, and cells were washed with PBS prior to lysis in RIPA buffer containing protease and phosphatase inhibitors. Protein concentration was determined by BCA Assay prior to western blot analyses.

### Statistics

Statistical tests and graphical representation achieved in GraphPad Prism.

## ACKNOWLEDGMENTS

This study was supported by grants from the National Institutes of Health to SML (GM146769) and MS (AI170877), as well as from the American Lung Association to SML.

## AUTHOR CONTRIBUTIONS

SML conceptualized the study. RJ, AB, NK, AG, MG, SG, AA, SML performed experiments. SML, RJ, AB, and MB analyzed the data. MS supervised the virus work. SML wrote the manuscript with contributions from all authors.

---

**Supplementary Figure 1.**
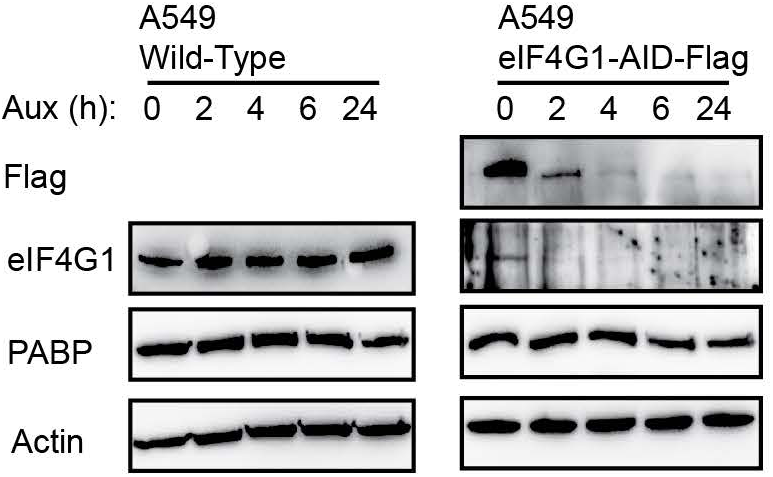

**Supplementary Figure 2.**
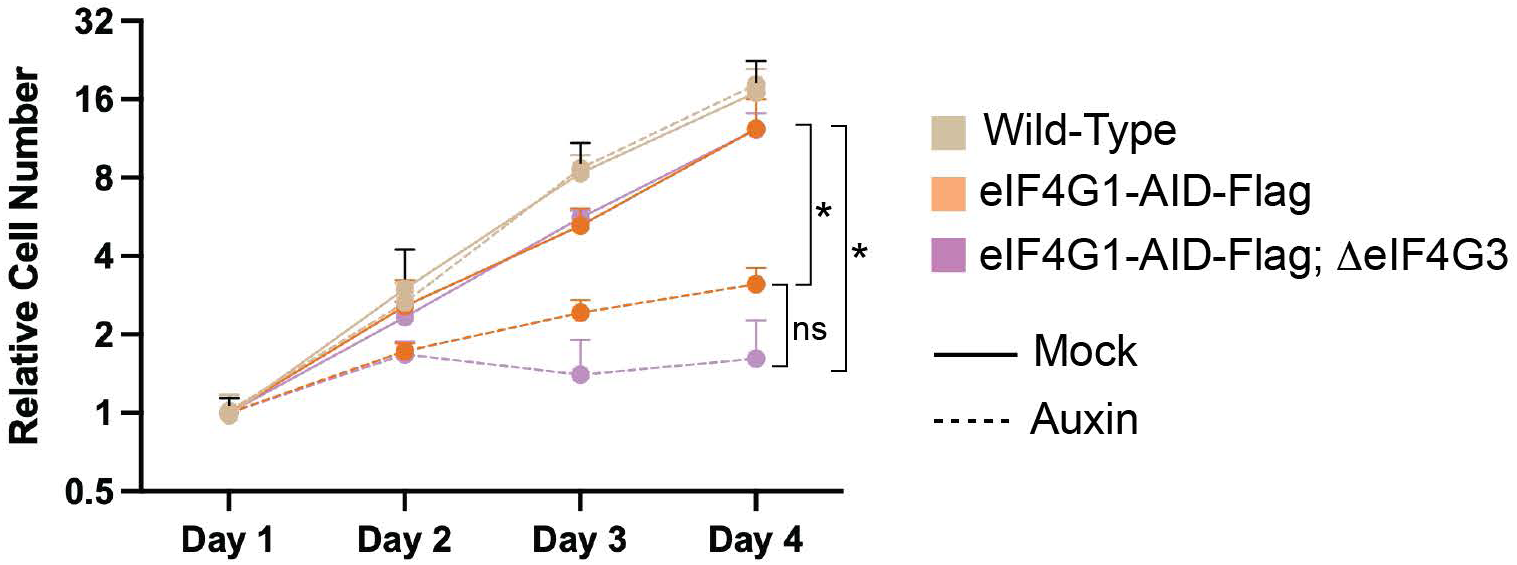

**Supplementary Figure 1.**
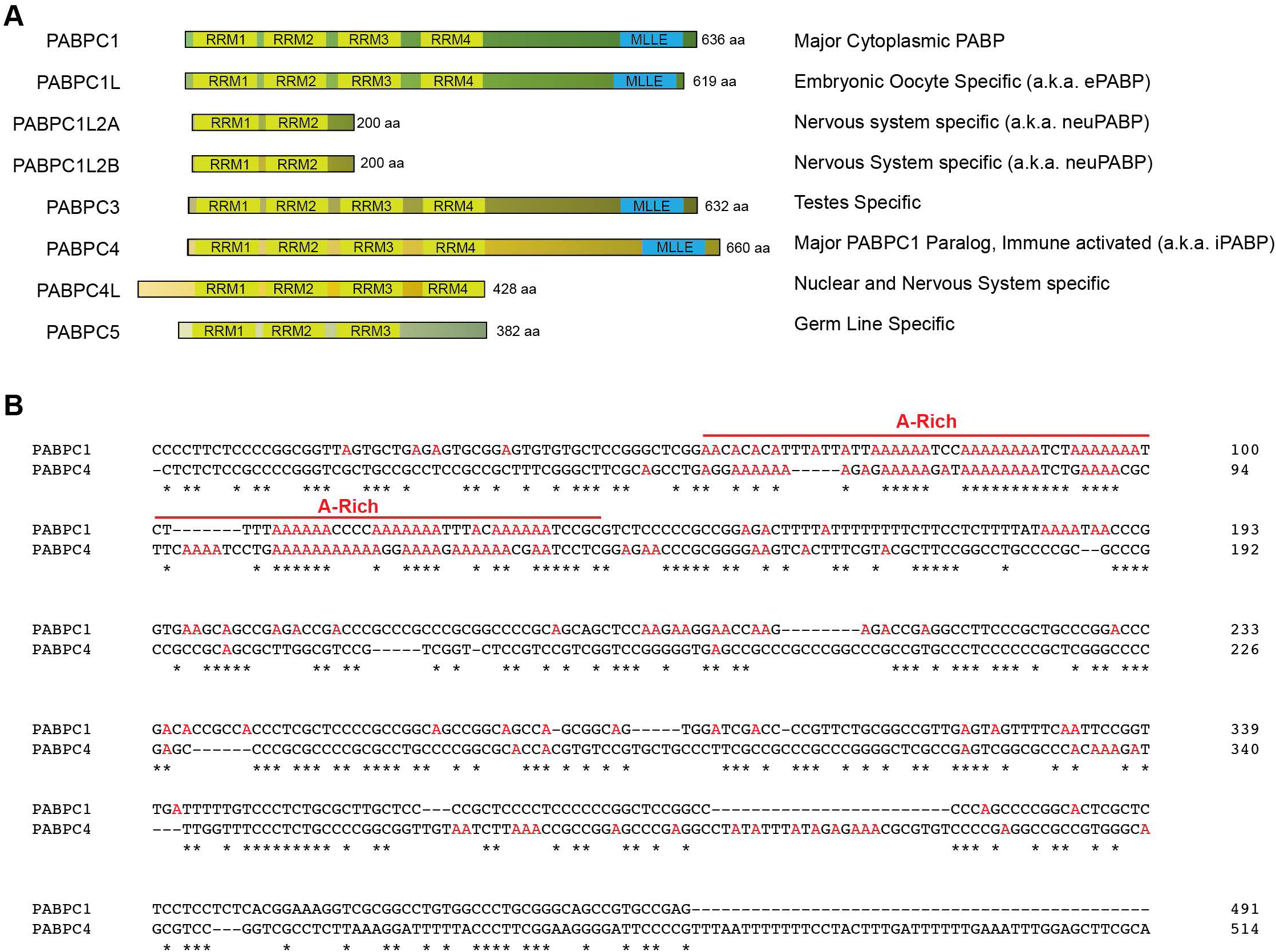

